# AbTune: Layer-wise Selective Fine-tuning of Protein Language Models for Antibodies

**DOI:** 10.1101/2025.10.17.682998

**Authors:** Xiaotong Xu, Alexandre M.J.J. Bonvin

## Abstract

**Motivation:** Antibodies play central roles in immune defense and are widely used as therapeutic agents. However, the high structural and sequence diversity of antigen-binding loops, combined with limited experimental data and weak co-evolutionary signals, makes it difficult to develop generalizable predictive models.

**Results:** We investigate test-time fine-tuning strategies to improve protein language model (pLM) performance in low-data settings, with a focus on antibody-related tasks. Systematic evaluations show that carefully constrained fine-tuning improves performance while preserving generalization. In particular, depth-selective fine-tuning consistently outperforms full-depth fine-tuning, with optimal performance achieved when tuning 50–75% of model layers for medium- to small-sized pLMs. We introduce **AbTune**, a test-time fine-tuning framework leveraging this depth-controlled adaptation strategy. Across antibody structure prediction, mutation effect prediction, and binding affinity prediction, AbTune outperforms standard pLM baselines and task-specific predictors on most benchmarks. We further analyze representation shifts, sequence-dependent adaptation behavior, and overfitting indicators, showing that fine-tuning depth, duration, and perplexity jointly determine performance.

**Availability:** https://github.com/haddocking/AbTune

## 1 Introduction

Antibodies are essential components of our immune system. They recognize and bind specific antigens to neutralize pathogens. Structurally, antibodies are Y-shaped molecules composed of two identical heavy and light chains. At the tips of each antibody arm lie the variable domains, which contain the complementarity-determining regions (CDRs) responsible for antigen recognition. Each antibody contains six CDR loops, three on the heavy chain and three on the light chain. The diversity of CDRs both in length, sequence and structures underlies the adaptability of antibodies, allowing them to recognize a vast array of antigens. Among these, the third heavy chain CDR loop (CDR H3) shows the largest sequence variability and structural diversity, and often contributes the majority of contacts with the target epitope [1].

This diversity in the CDR regions, combined with the lack of evolutionary signal for these loops, makes it challenging to develop generalizable predictors for any antibody-related tasks. For example, accurately predicting structures of antibodies and their complexes remains difficult. Although the state-of-the-art co-folding algorithm AlphaFold3 [2] has improved performance on such complexes through architectural innovations, successful predictions still require extensive sampling and accurate scoring. As a result, integrative modeling approaches based on docking or simulations remain highly valuable for these systems [3, 4]. Further challenges arise in predicting the binding affinity of antibody–antigen complexes and the effects of mutations on binding affinity. Recent studies show that current predictors generalize poorly in ΔΔ*G* prediction, and that future progress is fundamentally constrained by the limited quantity and variability of experimental data [5].

Protein Language Models (pLMs) have emerged as a powerful class of foundation models with broad applicability across diverse biological tasks [6–8]. The increasing availability of curated antibody sequence datasets, such as the Observed Antibody Space (OAS) [9], has further enabled the development of antibody-specific LMs. Early models that trained from scratch on OAS including AbLang [10] and AntiBERTy [11], demonstrated strong performance in tasks such as sequence recovery and paratope prediction. More recent models [12] leverage paired heavy and light chain sequences to capture cross-chain dependencies. Such domain-specific or task-specific adaption has been shown to be always beneficial [13, 14]. However, fine-tuning antibody-specific LM remains computationally expensive and data hungry, even when parameter-efficient fine-tuning (PEFT) techniques [15, 16] are applied.

In this work, we present AbTune, an algorithm for investigating the extent to which pLMs can be leveraged for antibody-related applications in the absence of additional training data, relying on selective test-time finetuning. Our main contributions are three-fold. First, we extended test-time fine-tuning strategies [17] to three biologically relevant antibody applications and provide further evidence that this approach yields consistent performance gains across multiple tasks. Furthermore, we systematically investigate the impact of fine-tuning depth on model performance and uncover that the optimal depth may depend on the size of the language model. In addition, we examine how embeddings evolve during fine-tuning and investigate when and how over-fitting potentially arises in this setting. Finally, using our AbTune protocol, we achieve state-of-the-art performance on two applications, highlighting its utility for computational immunology.

## 2 Methods

### 2.1 ESM Models

ESM (Evolutionary scale Modeling) [6, 8] is a family of transformer-based protein language models (pLMs) that use self-attention mechanisms to capture contextual relationships between amino acid residues in protein sequences. Given an input sequence, the model generates contextual embeddings by iteratively refining token representations across multiple transformer layers. ESMFold builds upon ESM-2 embeddings for protein structure prediction by first processing sequence embeddings through a Folding Trunk module that integrates contextual information across the sequence, followed by a Structure Module that predicts 3D atomic coordinates.

In this work, we studied four ESM model variants with parameter sizes ranging from 8 million to 3 billion: t6, t12, t30 and t33.

### 2.2 layer-wise selective LoRA fine-tuning

We extended the test-time fine-tuning approach originally proposed by [17]. Given a sequence input, pLM is fine-tuned using a masked language modeling (MLM) objective, in which random tokens are masked and predicted. Formally, the MLM loss is defined as:

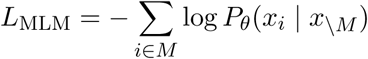

where *x* denotes the input sequence tokens, *M* the set of masked positions, and *θ* the model parameters. Optimization is carried out with stochastic gradient descent (SGD) using zero momentum and weight decay.

To enable efficient fine-tuning of large pLMs such as ESM, we employed Low-Rank Adaptation (LoRA) [15] by updating only the Linear layers within the Multi-Head Attention modules. Specifically, the weight update is defined as:

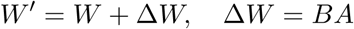

where *W* are frozen pretrained weights, and *A* ∈ *R^r^*^×*d*^*, B* ∈ *R^d^*^×*r*^ are trainable low-rank matrices with rank *r* = 4 and scaling factor *α* = 32. Following prior work [18], we further explored layer-wise selective fine-tuning by updating only a subset of LoRA layers. Denoting the full parameter set as Θ, selective fine-tuning can be expressed as:

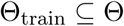

where Θ_train_ corresponds to the subset of layers chosen (25%, 50%, 75%, or 100%). All fine-tuning experiments were implemented with AbTune, which is available at https://github.com/haddocking/AbTune. The training procedure is highly efficient, although runtime varies depending on factors such as sequence length, ESM model size, fine-tuning depth, and the number of training epochs. For example, on our local cluster equipped with NVIDIA RTX A6000 Ada GPUs, fine-tuning a 113-residue sequence using the esm2 t33 650M UR50D model for 10 epochs took 1.14 seconds.

### 2.3 Perplexity Calculation

In pLMs, perplexity captures the model’s average ability to predict residues at each position in a sequence, and therefore reflects how well the model assigns probability to a given protein sequence. It ranges from 1 to infinity, with lower values indicating better predictive performance. We adopt the definition of perplexity from [17]:

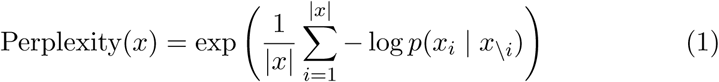

where |*x*| is the length of the protein sequence, and *p*(*x_i_*| *x*_\_*_i_*) is the probability that the model correctly predicts the residue *x_i_* at position *i* when it is masked.

### 2.4 Application 1: Antibody structure prediction

#### 2.4.1 Dataset: SAbDab structure dataset

Antibody structure prediction remains a challenging problem in computational biology, especially for CDR regions. This task aims to evaluate whether the proposed fine-tuning scheme can produce improved embeddings that lead to better structure predictions.

To construct the dataset for antibody structure prediction task, we downloaded the full SAbDab database [19](April 2025 release). We then applied the following filters: (1) X-ray crystal structure with resolution better than 3.5Å, (2) all CDR loops should contain more than two residues, (3) 100% similarity filtering on the CDR sequence, such that when multiple antibodies have identical CDR sequences, only one structure is kept, (4) The structures should have been released after May 2020. After applying these filters, the final benchmark set consists of 763 antibody structures.

#### 2.4.2 Method: Fine-tuning ESMFold for improved structure prediction

We used a local version of ESMFold (ESMFold-v1) to predict antibody structures. To minimize the effect of chain packing on the calculation, we first separated and aligned the heavy and light chains of each antibody, and then computed the RMSD for each loop separately. The chain-packing capability of ESMFold was evaluated by aligning the full predicted structure to the experimental structure and applying the CAPRI criteria [20] for acceptable models (Fnat ≥ 0.10, iRMSD ≤ 4.0 Å, and lRMSD ≤ 10.0 Å) to compute the success rate.

In the fine-tuning pipeline, we first extracted antibody sequences from our benchmark dataset. For each antibody, the heavy and light chain sequences were concatenated using a 25-residue polyglycine linker, following the strategy employed by ESMFold for complex prediction. The resulting full-length sequences were then used as inputs for fine-tuning. We focused our fine-tuning efforts on the protein language model esm2 t36 3B UR50D, which serves as the backbone language model of ESMFold (v1). Unlike prior fine-tuning strategies that focus on the Folding Trunk [21], we fine-tuned only the linear projection layers within the Multi-Head Attention modules of ESM-2, while keeping all parameters in the Folding Trunk module frozen. Four fine-tuning depths were evaluated: 25%, 50%, 75%, and 100%. Each sequence in our dataset was fine-tuned for 50 steps, during which we saved the updated embeddings and the corresponding antibody structures predicted by ESMFold at each step. The linker was then removed, RMSD for each region as well as chain-packing success rates were computed. In the final results, we reported the best values observed over the 50 fine-tuning steps.

We further analyzed the evolution of ESM embeddings during fine-tuning both qualitatively and quantitatively. To visualize a typical fine-tuning process, we applied UMAP to project embeddings obtained during fine-tuning of a representative antibody (PDB ID: 7mzn) into a two-dimensional space defined by real protein sequences from the Astral40 dataset and an equal number of random sequences from Astral40R [22], which consist of randomly shuffling sequences from Astral40. To quantify embedding dynamics, we computed the nearest-neighbor Astral40 distance (NAR) throughout finetuning to assess embedding reliability. Specifically, for each antibody, we measured the distance in embedding space to its nearest neighbor in the Astral40 reference set at each fine-tuning step. We then tracked NAR values across all antibodies in the benchmark over 50 fine-tuning steps. Based on their temporal behavior, antibodies were categorized into two classes: stable and drifting, corresponding respectively to embeddings that remain close to or progressively diverge from the Astral40 manifold over the course of fine-tuning.

### 2.5 Application 2: Zero-shot prediction of beneficial mutations

#### 2.5.1 Dataset: Antibody Point Mutation Dataset

Our dataset consists of 47 antibody–antigen complexes drawn from three sources: SKEMPI v2 [23, 24], AB-Bind [25], and AbDesign [26]. It contains a total of 1,303 point mutations with experimentally measured ΔΔ*G* values, including 49.4% affinity-improving mutations and 50.5% neutral or detrimental mutations, spanning different CDR regions. The distribution of the number of mutations per complex is available in SI Figure 1.

**Figure 1:**
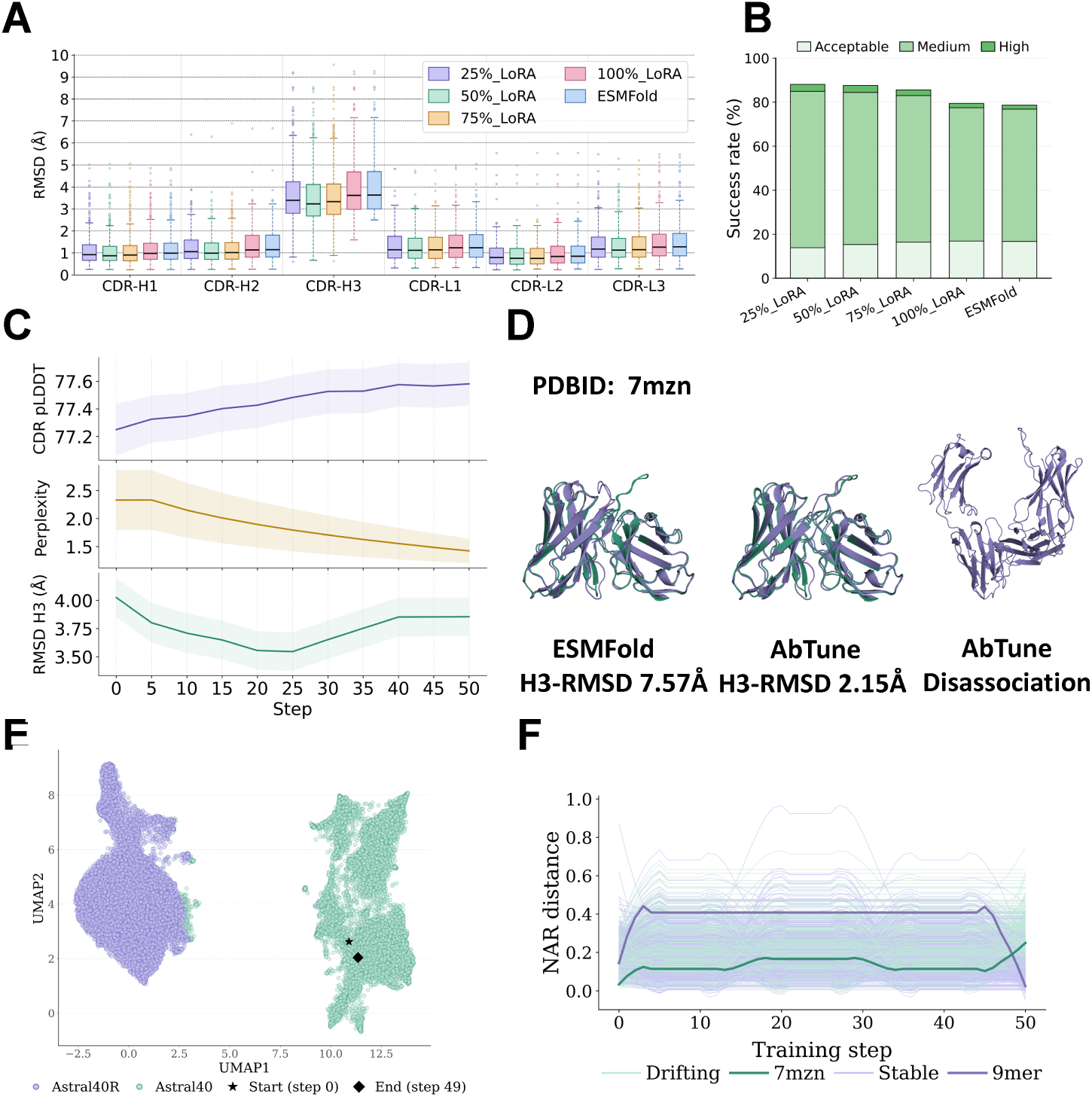
Antibody structure prediction performance and fine-tuning dynamics.**(A)** Distributions of RMSD across the six antibody CDR loops for five methods in our benchmark. ESMFold refers to the baseline ESMFold (v1) model without fine-tuning. Labels ‘XX%_LoRA_’ indicate AbTune finetuning where ‘XX%’ denotes depth of LoRA layers fine-tuned. RMSD values are computed from the best-performing fine-tuning step among 50 steps. **(B)** Heavy–light chain packing performance evaluated using CAPRI criteria across the five methods described above. **(C)** Dynamics of H3 RMSD, CDR pLDDT, and sequence perplexity during fine-tuning, shown in three panels with a shared x-axis of fine-tuning steps. Curves denote mean values across all sequences, with shaded regions indicating standard deviations for better visualization. **(D)** Representative antibody structure (PDB ID: 7MZN) showing substantial improvement with AbTune: H3 RMSD decreases from 7.57 Å to 2.15 Å. Chain disassociation was observed at the end of fine-tuning. **(E)** UMAP visualization of ESM em^1^b^8^eddings for a representative antibody (PDB ID: 7MZN) before and after fine-tuning. **(F)** Nearest-neighbor Astral40 distance (NAR) analysis in embedding space during fine-tuning. Antibodies are categorized into two behavioral classes, stable and drifting, with representative examples highlighted in darker colors (PDB ID: 7MZN for drifting; PDB ID: 9MER for stable).

The ΔΔ*G* values are reported in two formats: either directly as the change in binding free energy (ΔΔ*G*) after mutation, or as the ratio of ELISA affinity between the wild-type and mutant. A positive or zero ΔΔ*G*, or a ratio greater than or equal to 1, indicates weaker binding, and such mutations were labeled 0 in our dataset. We treated the prediction task as binary: mutations predicted to improve binding were labeled 1 (beneficial), and all others were labeled 0 (non-beneficial).

#### 2.5.2 Method: Fine-tuning ESM for improved prediction of beneficial mutations

For both the baseline and fine-tuned ESM models, we use predicted probabilities as a proxy for binding affinity to assess the impact of mutations. Given a protein sequence with a mutation at position *i*, the residue at that position is masked and passed through the model. The resulting logits are then normalized into probability distributions using the softmax function:

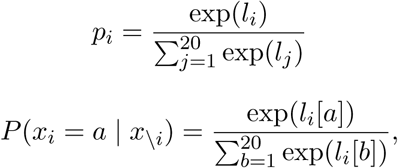

where *l_i_*[*a*] denotes the logit corresponding to amino acid *a* at position *i*, and *x*_\_*_i_* represents the sequence with position *i* masked. From this distribution, we extract the probabilities corresponding to the wild-type (*P*_WT_) and mutant (*P*_MUT_) residues. If *P*_WT_ *< P*_MUT_, the mutation is considered beneficial for binding and labeled as 1; otherwise, it is labeled as 0.

We fine-tuned four ESM model variants including t6, t12, t30, and t33. Each model was fine-tuned using LoRA at four different depths: 25%, 50%, 75%, and 100%, resulting in 16 fine-tuning configurations. For each wild-type antibody–antigen complex in the benchmark, the antibody heavy chain, light chain, and antigen sequences were concatenated into a single joint sequence as input.

For every antibody-antigen complex, mutation effects were first predicted using the pretrained baseline models (ESM models without fine-tuning) and evaluated using Accuracy, Precision, F1 and MCC score. The results were then averaged across all complexes. All ESM models were then fine-tuned for 50 steps on every wild-type sequence, and predictions were re-evaluated with the same metrics. The best-performing AbTune configuration for each ESM model is reported in Figure 2A. As external baselines, we evaluated AbLang2 and AntiBERTy using only the combined heavy and light chain sequences of the antibody, without antigen information. For MSA-based language models, paired MSAs for antibody–antigen complexes were generated using MMseqs2 [27], following the approach described in MSA Pairformer [28]. We ran RDE-PPI [29] using the default settings and used the pdbs aligned structures prepared by Janusz et al. [26] as wild-type input structures. Three VESM models [30] were also evaluated using the code and released model checkpoints provided by the authors.

**Figure 2:**
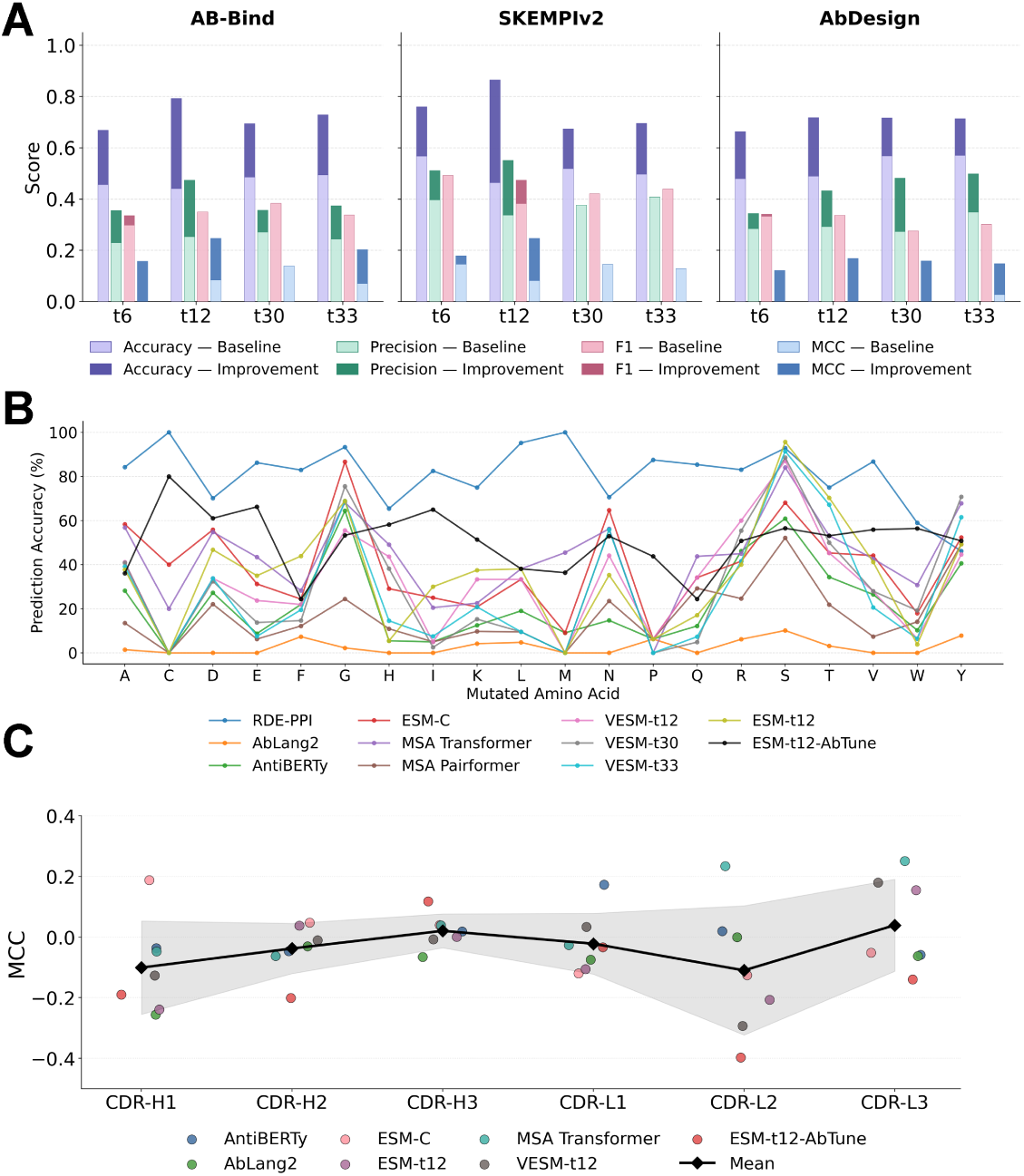

### 2.6 Application 3: Binding Affinity Prediction

#### 2.6.1 Dataset: Observed Antibody Space (OAS) dataset

The OAS database contains annotated large-scale immune repertoires, encompassing over one billion sequences across diverse immune states in both human and mouse subjects [9]. To estimate the binding affinity, we use sequence redundancy as a proxy as established in previous works [31, 32]. Antibodies that bind strongly to their targets are preferentially selected and clonally expanded, resulting in their sequences being observed more frequently in the repertoire.

To train and evaluate our binding affinity predictors, we downloaded and processed all paired sequences following the detailed protocol described in AntiFormer [31]. Our final dataset comprised 1,476,057 paired antibody heavy and light sequences, each assigned a label of 1 or 0, with 15.7% labeled as 1 (high-affinity) and 84.3% labeled as 0 (low-affinity). The dataset was randomly partitioned into five subsets for 5-fold cross-validation.

#### 2.6.2 Dataset: Antibody chain pairing test set

We extend our analysis to the task of predicting correct heavy–light chain pairing. Three independent paired VH–VL datasets from ImmunoMatch were used for evaluation [14], including tonsil B-cell repertoires [33], leukemia and lymphoma samples from the Cancer Cell Line Encyclopedia (CCLE) [34], and chronic lymphocytic leukaemia (CLL) cohorts [35]. For each dataset, we constructed an equal number of negative samples by randomly shuffling true VH–VL pairs. The performance of both ImmunoMatch [14] and BindFormer-v3 was assessed under a ranking-based evaluation protocol as suggested. Specifically, for each VH–VL candidate pair, the one with the higher predicted score was selected as the predicted true pairing and classification metrics for this task were reported.

#### 2.6.3 Method: Fine-tuned Embeddings for Improved Prediction of Binding Affinity

We used ESM-2 embeddings from the model esm2 t33 650M UR50D as features for each antibody pair (heavy and light chains). Sequence-level representations were obtained by averaging across the embedding dimension. This model was chosen as it provides a practical balance between computational efficiency and performance. The averaged embeddings were then padded to the length of the longest chain in the dataset.

We developed BindFormer, a lightweight dual-chain classifier for binding affinity prediction. Each chain was independently encoded using a rotary multi-head self-attention (RoPE-MHA) module [36], which captures contextual dependencies while encoding relative positional information. The representations were reduced to fixed-length embeddings via attention pooling. The heavy- and light-chain embeddings were then concatenated and passed through a MLP classification head to generate the final prediction. The model architecture and training details are described in SI Method section.

As a baseline, each fold was trained using embeddings directly from ESM-2 without any additional fine-tuning. Final performance metrics were averaged across the five folds as in AntiFormer [31]. Given the computational cost of fine-tuning on every sequence in OAS, we adopted a rank-based partial fine-tuning strategy. Sequences were ranked by perplexity, and only the top-ranked sequences were fine-tuned for 50 steps as sequences in this dataset generally have higher starting perplexity. Based on prior experience with fine-tuning larger ESM models, 50% of the LoRA layers were finetuned and BindFormer was then retrained using the fine-tuned embeddings. Fine-tuning was performed on the top 0.1% and 1% of sequences to evaluate performance gains as well as a random 1% subset. Reported performances of other methods (AntiFormer, AntiBERTy, LlamaAffinity) were taken from corresponding publications [31, 32]. All BindFormer models were trained for 75 epochs on a single NVIDIA RTX A6000 Ada GPU. We used the AdamW optimizer with a learning rate of 1 × 10^−4^, weight decay of 1 × 10^−4^, and a batch size of 128, along with a Cosine Annealing learning rate scheduler and cross-entropy loss.

## 3 Results

### 3.1 Improved antibody structure prediction

We first assessed the performance of ESMFold [6] on the task of antibody structure prediction using a challenging benchmark comprising 763 antibodies (See Methods for detail). Overall, ESMFold struggles to generalize to antibodies outside its training distribution (Figure 1A), yielding substantially higher RMSD values than those reported for AlphaFold2 [3], and AlphaFold3 [37]. Consistent with previous studies, the H3 loop remains the most challenging region to predict accurately. In addition, analysis shows that ESMFold frequently fails to predict the relative orientation between the heavy and light chains: in 24.1% of cases, the output antibody structures do not satisfy CAPRI criteria for acceptable models (Figure 1B).

To address these limitations, we next investigated whether fine-tuning can improve the prediction capabilities of ESMFold. As shown in Figure 1A, fine-tuning consistently improves antibody structure predictions across all CDR regions, regardless of the proportion of LoRA layers fine-tuned. In particular, a mean of 11.9% reduction in H3 RMSD is achieved when finetuning the first 50% of LoRA layers. Beyond improvements in mean accuracy, the lower tail of the CDR-H3 RMSD distribution shifts significantly downward after fine-tuning, with four antibodies achieving sub-angstrom accuracy (¡1 Å), indicating that fine-tuning improves not only average performance but also enables highly accurate predictions for a subset of challenging targets. Heavy–light chain packing success increases from 78.6% (ESMFold baseline) to 87.5% with the best AbTune protocol, suggesting that joint fine-tuning of heavy and light chains improves inter-chain contact prediction in ESMFold, in line with the reported benefits of supplementing antibody LMs with native chain pairing [38].

For both sub-tasks, the best performance is achieved when only the first 50% of layers are fine-tuned, whereas fine-tuning all layers leads to worse performance than partial fine-tuning, in line with previous evidence [18]. To further explore the learning dynamics of pLMs, we tracked how H3 RMSD, CDR pLDDT, and sequence perplexity evolved at each fine-tuning step for all sequences in our dataset (Figure 1C). In contrast to the strong correlation between TM-score and pLDDT reported for general proteins [17], we found that the fine-tuning step producing antibody structures with the lowest RMSD does not always correspond to the highest pLDDT or lowest sequence perplexity.

As shown in recent benchmarks of AlphaFold3 performance on antibodies [39, 40], confidence metrics produced by co-folding models, while useful, are not always reliable predictors of structural accuracy. The same holds for ESMFold. With respect to sequence perplexity, Pugh *et al.* [41] showed that its relationship with biological fitness is not straightforward, as it is influenced by training data biases and phylogenetic relationships among sequences. Consistent with these observations, we find that ESMFold is less reliable when the corresponding perplexity from ESM-2 lies at either extreme—either too high or too low [42, 43].

A representative antibody structure from our benchmark (PDB ID: 7mzn) is shown in Figure 1D. AbTune reduces H3 RMSD by 71.6%, from 7.57 Å (ESMFold) to 2.15 Å. Interestingly, after 50 fine-tuning steps, the antibody begins to dissociate despite the presence of a linker connecting the two chains. To gain more insights into the fine-tuning process and identify possible explanations for this behavior, we analyzed the evolution of embeddings. Inspired by Random Neighbor Score (RNS) [22] as a quality measurement for pLM embeddings, we projected the embeddings of the representative antibody at each fine-tuning step into the reference embedding space defined by the Astral40 and Astral40R (R stands for random) protein sequence sets (Figure 1E). As seen in the Figure, the Astral40 space is densely packed, with the initial embedding of 7mzn situated near the center of the distribution. Comparing embeddings obtained at the starting and final steps suggests that the overall displacement is relatively small, while remaining within the defined space of natural proteins in Astral40. Interestingly, despite only small changes in the embedding space, substantial improvements in structure prediction performance were observed, as described above.

Quantitative analysis of the distance to the Nearest Astral40 Representative (NAR) across fine-tuning steps (Figure 1F) for antibodies in our dataset reveals two distinct modes: Stable and Drifting. Stable fine-tuning remains close to the Astral40 manifold with minimal changes in distance and k-nearest neighbor composition, whereas drifting shows increased distance during fine-tuning without recovery, indicating persistent displacement in embedding space. The trajectory of 7mzn highlights this behavior: the NAR distance rises sharply in early steps, stabilizes at an intermediate plateau between steps 18 and 20, and then increases abruptly toward the final steps ( 50). This may explain antibody dissociation described earlier at the end of fine-tuning. Importantly, across all complexes in the dataset, the magnitude of the observed distance changes is generally small. This indicates that finetuning primarily involves localized exploration of weight space to identify optimal parameters for a given sequence, resulting in only modest shifts in the embedding manifold. Examining all distance trajectories in Figure 1F, together with signs of structural degradation under prolonged fine-tuning, we conclude that the potential window in which fine-tuning is beneficial is both narrow and highly sequence dependent. In particular, when the initial perplexity is low, a small number of fine-tuning steps is sufficient to achieve performance gains, whereas extended fine-tuning can be detrimental.

Benchmarking beneficial mutation prediction. **(A)** Performance comparison of sequence-based models on beneficial mutation prediction across three benchmark datasets: AB-Bind, SKEMPIv2, and AbDesign. The figure reports Accuracy, Precision, F1, and MCC for multiple model variants, shown before and after fine-tuning with LoRA as stacked bars. The best fine-tuning depths for ESM model t6, t12, t30, and t33 are 75%, 75%, 50%, and 50%, respectively. For each dataset, the metrics are reported at the best fine-tuning step. **(B)** Distribution of predicted beneficial mutations across amino acid types. The x-axis shows the 20 standard amino acids, and the y-axis indicates the percentage of predictions for which a mutation to that amino acid was classified as beneficial (predicted label = 1) by each model. **(C)** MCC values across antibody CDR regions. Individual points denote the performance of each model, and the black line highlights the mean across methods.

### 3.2 Improved zero-shot prediction of beneficial mutations of antibody-antigen complexes

Identifying point mutations that enhance antibody–antigen binding is a crucial step in the development of therapeutic antibodies once an initial binder has been identified. However, this task remains extremely challenging due to the complexity of molecular interactions and the limited availability of experimental data in this regime. Previous work [26] has shown that three state-of-the-art structure-based predictors struggle to generalize to antibody–antigen complexes outside their training distributions. Moreover, stratification by mutation type reveals substantial dataset and model biases; for example, tyrosine is the most frequently mutated residue in experiments, while alanine substitutions are most common due to alanine scanning protocols [26].

Considering the biological relevance and difficulty for this task [5], we selected prediction of beneficial mutations in antibody–antigen complexes as the second AbTune application. We formulated the problem as a binary classification task: given a point mutation in a mutated antibody-antigen complex, the goal is to predict whether the mutation is beneficial or nonbeneficial (neutral or detrimental) towards binding as measured by ΔΔ*G*. This formulation was chosen instead of regression because the available dataset contains heterogeneous ΔΔ*G* measurements and is highly imbalanced across complexes, making classification a more robust alternative (SI Figure 1).

We first evaluated the zero-shot performance of four ESM variants of increasing scale using probability outputs as predicted binding affinities. Model performance was assessed on three datasets: SKEMPIv2 [23, 24], AB-Bind [25], and AbDesign [26], using four standard classification metrics. As shown in Figure 2A, both AbDesign and AB-Bind are consistently more challenging than SKEMPIv2 across all models. Furthermore, the performance of ESM models scaled poorly with model size, a phenomenon previously reported in several studies [44, 45].

To assess the impact of AbTune, all four ESM variants were fine-tuned using four different depths, defined by the proportion of layers fine-tuned via LoRA. Across all models and datasets, fine-tuning consistently improved the performance significantly (Figure 2A). Consistent with observations in structure prediction tasks, optimal performance was achieved by fine-tuning only a subset of layers depending on ESM model scale: 75% for small to medium-sized models and 50% for larger models. Across all four finetuning settings, a substantial improvement in Matthews correlation coefficient (MCC) was observed, corresponding to an average relative increase of more than tenfold, indicating that fine-tuning extends beyond simple memorization of wild-type sequences. We also tested whether sequence-specific properties may inform optimal AbTune configurations. A moderate correlation (r = 0.585, SI Figure 2) was observed between the initial perplexity of a sequence and the number of fine-tuning steps required to reach optimal performance. This suggests that input sequence characteristics may serve as a practical indicator for determining the required fine-tuning duration across sequences.

To provide a comprehensive benchmark under the considered problem setting, in Table 1 we evaluated ten other predictors: one structure-based predictor (RDE-PPI [29]), two antibody-specific pLMs (AbLang2 [46], AntiBERTy [11]), two from recent member of the ESM family [47], three variant-effect ESM models (VESM) models from ESM co-distillation [30], and two MSA-based pLM (MSA-Transformer [48], MSA Pairformer [28]). RDE-PPI, despite being trained on labeled ΔΔ*G* data and demonstrating strong performance on the SKEMPIv2 dataset, showed very poor generalization to the AB-Bind and AbDesign datasets. Since both AbLang2 and AntiBERTy are antibody-specific pLMs, predictions from these models are made without considering the antigen. we hypothesize that they primarily capture the effects of mutations on antibody stability or folding rather than binding. ESM-C, promoted as superior to ESM-2, indeed had improved performance on this task. MSA-based pLMs have demonstrated some advantages over single-sequence pLMs across various tasks [28, 48]. In our experiments, however, these benefits were observed primarily for MSA Transformer [48], but not for the more recent MSA Pairformer [28]. VESM also performed poorly on this dataset; however, it exhibited better scaling capabilities compared to the original ESM model. Among all external predictors evaluated, no single method consistently outperformed the others, with most MCC values remaining close to zero. The best performance was obtained by our fine-tuning protocol, ESM-t12-AbTune, which achieved a mean MCC of 0.228.

**Table 1:** Performance of sequence- and structure-based models on beneficial mu<u>tation prediction.</u>.

| Model | Accuracy | Precision | Recall | F1 | MCC |
| --- | --- | --- | --- | --- | --- |
| ESM-t12-AbTune | 0.393 | **0.404** | 0.506 | **0.814** | **0.228** |
| esm-t12 | 0.467 | 0.305 | 0.628 | 0.361 | 0.053 |
| VESM-t12 | 0.494 | 0.314 | 0.548 | 0.361 | 0.027 |
| VESM-t30 | 0.503 | 0.308 | 0.580 | 0.362 | 0.024 |
| VESM-t33 | 0.527 | 0.348 | **0.630** | 0.399 | 0.128 |
| ESMC-300 | 0.517 | 0.282 | 0.439 | 0.312 | -0.015 |
| ESMC-600 | 0.508 | 0.298 | 0.428 | 0.314 | -0.033 |
| MSA Pairformer | 0.580 | 0.201 | 0.168 | 0.154 | -0.080 |
| MSA Transformer | 0.494 | 0.285 | 0.475 | 0.315 | -0.018 |
| AbLang2 | 0.693 | 0.117 | 0.037 | 0.055 | -0.006 |
| AntiBERTy | 0.557 | 0.243 | 0.293 | 0.228 | -0.061 |
| RDE-PPI | **0.713** | 0.347 | 0.228 | 0.228 | 0.146 |

To investigate whether dataset biases influenced model predictions, we analyzed each amino acid substitution (i.e., mutations to a given target residue) by computing the proportion of cases in which each model predicts the mutation to be beneficial for binding. As shown in Figure 2B, RDE-PPI exhibits a generally higher tendency to classify any mutations as beneficial compared to other methods. Although AbLang2 and AntiBERTy are both antibody-specific LMs, they showed distinct amino acid preference profiles, yet both showed a pronounced bias toward serine (S). Tyrosine (Y), which is among the most favored substitutions predicted by ESM-t12-AbTune and is also frequently preferred by other sequence-based models. This observation is consistent with reported antibody sequence statistics, which show a notable enrichment of Tyrosine residues in antibodies [49].

To further explore whether predictive performance varies across antibody regions, we stratified mutations by CDR location and evaluated model performance using MCC within each region. Surprisingly, as shown in Figure 2C, for AbTune, mutations in the CDR-H3 loop were, on average, easier to predict than those in other CDRs, with CDR-H3 accounting for 65.5% of all point mutations. Across all methods, however, no consistent pattern emerged, with different models exhibiting region-specific strengths and weaknesses.

### 3.3 Improved antibody binding affinity prediction

Predicting antibody binding affinity is critical for understanding immune responses and guiding therapeutic design. To address this challenge, we developed and trained an architecture named as BindFormer (see Methods). We formulate the problem as a binary classification task in which, given the heavy and light chain sequences of an antibody, the model predicts whether it is a binder or non-binder.

Our baseline model, which was trained directly on averaged ESM-2 embeddings without fine-tuning (BindFormer-ESM), already outperforms the current state-of-the-art method on this dataset AntiFormer [31] across all metrics except precision. In settings where large-scale fine-tuning may be required, we hypothesize that performance gains can be obtained by selectively fine-tuning sequences that are poorly represented in the pretrained embedding space, as quantified by perplexity.

To test this hypothesis, we first fine-tuned only the top 0.1% of sequences (BindFormer-v1), which already yielded consistent improvements across all metrics compared to the baseline. Expanding the fine-tuning set to the top 1% of sequences (BindFormer-v3) results in the best overall performance, outperforming four other predictors: AntiFormer [31], AntiBERTy [11], AntiBERTa [50], and the more recent LlamaAffinity [32] (Table 2). Fine-tuning a randomly selected 1% of sequences (BindFormer-v2) also improved performance, although not to the same extent as targeting the top 1% ranked by perplexity. Other fine-tuning settings were also explored by varying the number of fine-tuning steps; results are provided in SI Table 1.

**Table 2:** Performance comparison of BindFormer and existing predictors for antibody binding affinity. BindFormer variants include: BindFormer-esm (ESM embedding features without fine-tuning), BindFormer-v1 (top perplexity 0.1% fine-tuned), BindFormer-v2 (random 1% fine-tuned), and BindFormer-v3 (top perplexity 1% fine-tuned). Performance of other methods was taken from the original publications.

| Model | Accuracy | F1 | Precision | Recall | AUC |
| --- | --- | --- | --- | --- | --- |
| AntiFormer | 0.918 | 0.882 | 0.963 | 0.925 | 0.966 |
| AntiBERTy | 0.832 | 0.851 | 0.911 | 0.891 | 0.940 |
| LlamaAffinity | 0.964 | 0.964 | 0.970 | 0.959 | 0.994 |
| BindFormer-esm* | 0.942 | 0.943 | 0.945 | 0.942 | 0.980 |
| BindFormer-v1* | 0.956 | 0.951 | 0.957 | 0.956 | 0.990 |
| BindFormer-v2* | 0.973 | 0.973 | 0.973 | 0.973 | 0.995 |
| BindFormer-v3* | **0.977** | **0.976** | **0.977** | **0.977** | **0.996** |

\* denotes our method.

**Table 3:** Comparison of methods across chain pairing datasets (King Tonsil, CLL Sequences, and CCLE CellLines).

| Dataset | Method | Accuracy | Precision | F1 Score | MCC |
| --- | --- | --- | --- | --- | --- |
| King_Tonsil | ImmunoMatch | 0.501 | 0.501 | 0.500 | 0.002 |
|  | BindFormer-ESM | 0.499 | 0.503 | 0.351 | -0.001 |
|  | BindFormer-v3 | **0.524** | **0.524** | **0.523** | **0.048** |
| CLL_Sequences | ImmunoMatch | **0.525** | 0.525 | **0.525** | 0.050 |
|  | BindFormer-ESM | 0.513 | 0.516 | 0.483 | 0.030 |
|  | BindFormer-v3 | **0.525** | **0.536** | 0.486 | **0.060** |
| CCLE_CellLines | ImmunoMatch | 0.519 | 0.250 | 0.235 | -0.115 |
|  | BindFormer-ESM | 0.592 | 0.486 | **0.519** | -0.125 |
|  | BindFormer-v3 | **0.667** | **0.500** | 0.471 | **0.229** |

Beyond binding affinity, heavy–light (H–L) chain compatibility is a crucial determinant of functional antibody design, as proper pairing is essential for antibody stability and activity in both natural immune maturation and therapeutic engineering. However, predicting H–L pairing remains highly challenging. Our recent experience in CAPRI [20] also confirmed that even state-of-the-art models such as AlphaFold3 [2] can mis-pair heavy and light chains when multiple antibodies bind the same antigen. We therefore evaluated whether BindFormer can address this task, despite not being explicitly trained for chain pairing prediction.

We assessed three datasets of paired VH–VL sequences from ImmunoMatch [14], spanning diverse biological and disease contexts (see Methods for details). Despite not being trained for VH–VL pairing, BindFormer-v3 consistently improves over ImmunoMatch [14] across datasets, particularly in CCLE dataset where gains are more pronounced, despite ImmunoMatch being explicitly designed for this task. This suggests that fine-tuning helps capture meaningful signals underlying antibody chain compatibility. In contrast, BindFormer-ESM performs comparably to ImmunoMatch and did not exhibit the same performance improvements observed for BindFormer-v3 after fine-tuning.

We reviewed the literature to estimate the time and computational resources required for full training of antibody-specific LMs [10–12, 46, 51, 52]. While not all studies report training costs, available estimates suggest that training typically requires multiple days to several weeks and relies on distributed training across multiple high-performance GPUs.

In comparison, all fine-tuning experiments presented in this study required around 5 days of computation on our cluster equipped with 8 NVIDIA A6000 GPUs. A typical fine-tuning loop requires approximately 1 second per sequence on a single NVIDIA A6000 GPU. Even when scaled to the full OAS dataset [9], the total runtime would remain under one week, highlighting the computational efficiency of AbTune relative to other fine-tuning methods for antibody-specific LMs.

## 4 Discussion

In this work, we investigated whether test-time fine-tuning can serve as a practical mechanism for fine-tuning pLMs to antibody applications. We introduced AbTune, a lightweight sequence-specific, layer-wise fine-tuning method that improves performance across three biologically relevant tasks: antibody structure prediction, mutation effect prediction, and binding affinity prediction. Across these settings, AbTune consistently outperformed strong task-specific baselines and matched or exceeded prior best-performing methods on two of the three tasks.

Beyond overall gains, our results show that test-time fine-tuning is most effective when carefully constrained. Performance peaked when 50%–75% of LoRA layers were fine-tuned, suggesting a balance between retaining pretrained representations and adapting to each sequence. In most pLM applications, embeddings or logits are extracted from the last hidden layer [13, 53, 54]. However, information is not uniformly distributed across layers, and the last layer is not always optimal for downstream tasks [13, 55]. Furthermore, we found that optimal fine-tuning duration varies substantially across sequences and is partially correlated with initial sequence perplexity. This indicates that fine-tuning should be treated as a flexible process shaped by both model scale and input characteristics, rather than as a fixed protocol. Despite these improvements, important limitations remain. Identifying optimal AbTune configurations is non-trivial, performance is sensitive to hyperparameter choices, and overfitting is difficult to detect for non-structural tasks. While NAR-distance analysis can provide evidence in some cases, such signals are not always present. Moreover, mutation effect prediction and antibody chain pairing remain challenging, and it is still unclear to what extent language models capture biologically meaningful signals beyond evolutionary priors and germline bias [46, 56]. These limitations suggest several directions for improving robustness and explainability. Future work could focus on more robust hyperparameter selection and extend beyond single-sequence fine-tuning to mini-batch strategies. In particular, grouping sequences by properties such as CDR similarity may enable joint learning and better capture shared structural and biophysical patterns.

## Author contributions

Xiaotong Xu (Conceptualization [lead], Data curation [lead], Methodology [equal], Software [supporting], Visualization [lead], Writing–original draft [lead], Writing–review & editing [equal]), Alexandre M.J.J. Bonvin (Conceptualization [supporting], Data curation [supporting], Funding acquisition [lead], Methodology [supporting], Project administration [lead], Software [supporting], Supervision [lead], Writing–original draft [supporting], Writing–review & editing [equal]).

## Acknowledgments

This work was supported by the China Scholarship Council [202208310024 to X.Xu.].

## Data availability

All datasets used across the three applications are publicly available at https://doi.org/10.6084/m9.figshare.32351298. Source code and analysis pipelines are available at the official AbTune GitHub repository.

