## Supplementary material for "AbTune: Layer-wise Selective Fine-tuning of Protein Language Models for Antibodies"

SI Table 1. Performance of different BindFormer various under different fine-tuning scenarios.3

**SI Figure 1. Distribution of the number of data points per antibody-antigen complex for application 2** (mutation effect prediction)**.** Each bar represents the number of complexes with a given count of point mutation.

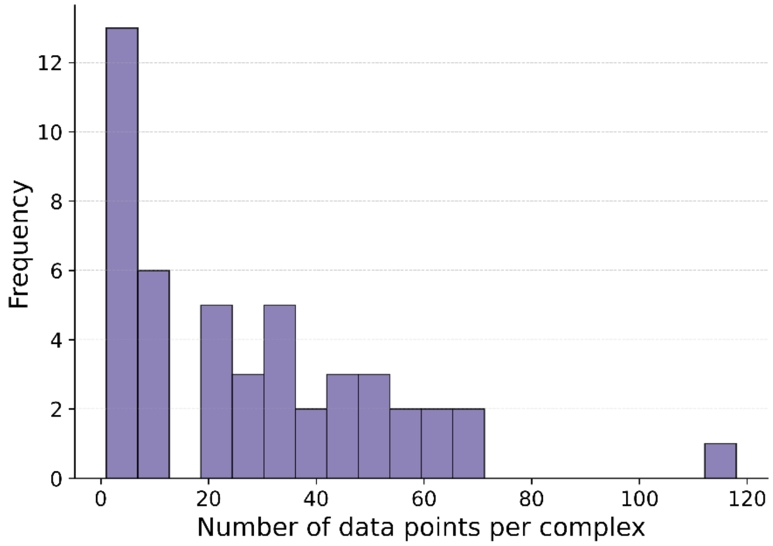

**SI Figure 2. The correlation between Perplexity and best performing step in application 2** (mutation effect prediction). We report here the moderate correlation between initial perplexity and the best forming step with our best performing model ESM-t12-AbTune.

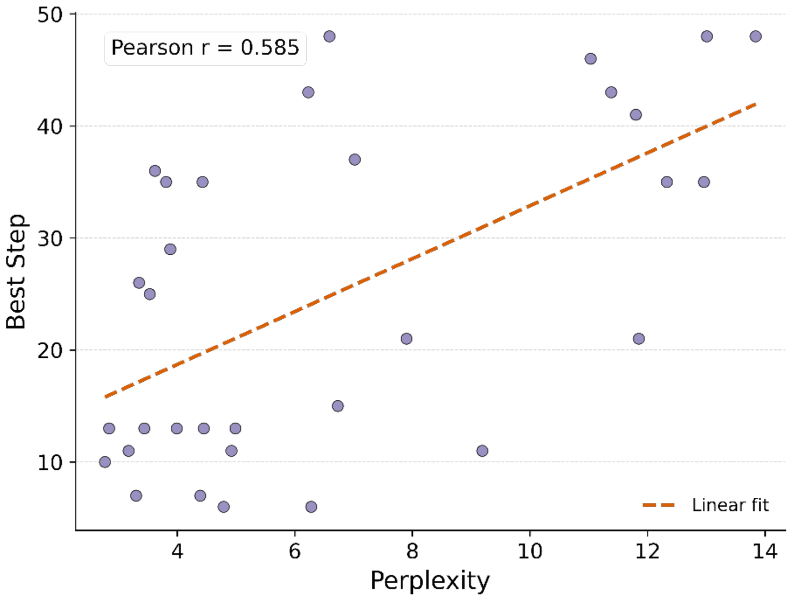

### SI Method. Training and model details of BindFormer

Our BinderFormer architecture models antibody heavy and light chain sequences using a dual-chain transformer-based encoder. Each chain is first embedded with ESM-2 (with default or fine-tuned weights) and then projected to a higher-dimensional space (d_attn = 128) via a linear layer. Sequential signals within each chain are captured using Rotary Multi-Head Attention (RoPE) with 4 attention heads. The attention outputs are then refined through a two-layer SiLU MLP with residual connections. A learned attention pooling module aggregates per-residue representations into a single vector for each chain. Finally, embeddings from the heavy and light chains are concatenated and passed through a three-layer fully connected classifier with ReLU activations, Batch Normalization, and Dropout (p = 0.2) for final classification output.

### SI Table 1. Performance of different BindFormer various under different fine-tuning scenarios

| Model (Vary by BinderFormer scenarios) | Accuracy | F1 | Precision | Recall | AUC |
|---|---|---|---|---|---|
| baseline | 0.942 | 0.943 | 0.945 | 0.942 | 0.98 |
| 0.1%-top-10step | 0.93 | 0.93 | 0.935 | 0.93 | 0.975 |
| 0.1%-top-50step | 0.956 | 0.951 | 0.957 | 0.956 | 0.99 |
| 0.1%-rand-10step | 0.951 | 0.952 | 0.953 | 0.951 | 0.982 |
| 0.1%-rand-50step | 0.977 | 0.977 | 0.977 | 0.977 | 0.996 |
| 1%-top-10step | 0.93 | 0.932 | 0.936 | 0.93 | 0.968 |
| 1%-top-50step | 0.977 | 0.976 | 0.977 | 0.977 | 0.996 |
| 1%-rand-10step | 0.906 | 0.878 | 0.855 | 0.906 | 0.789 |
| 1%-rand-50step | 0.973 | 0.973 | 0.973 | 0.973 | 0.995 |

Each model name follows the format <Percentage>% - <Selection Method> - <Training Steps>, where the percentage indicates the fraction of sequences selected for fine-tuning, the selection method specifies whether sequences were chosen based on highest perplexity(top) or randomly (rand), and the training steps indicate how many steps the model was fine-tuned on the selected sequences; for example, 0.1%-top-10step means the top 0.1% of sequences were fine-tuned for 10 steps.
